# Increase in primary cilia number and length upon VDAC1 depletion contributes to attenuated proliferation of cancer cells

**DOI:** 10.1101/2023.03.31.535181

**Authors:** Arpita Dutta, Priyadarshini Halder, Anakshi Gayen, Avik Mukherjee, Chandrama Mukherjee, Shubhra Majumder

## Abstract

Primary cilia (PCs) that are present in most human cells and perform sensory function or signal transduction are lost in many solid tumors. Previously, we identified VDAC1, best known to regulate mitochondrial bioenergetics, to negatively regulate ciliogenesis. Here, we show that downregulation of VDAC1 in pancreatic cancer-derived Panc1 and glioblastoma-derived U-87 cells significantly increased ciliation. Those PCs were remarkably longer than the control cells. Such increased ciliation inhibited cell cycle, which contributed to reduced proliferation of these cells. VDAC1-depletion also led to longer PCs in quiescent RPE1 cells. Therefore, serum-induced PC disassembly was slower in VDAC1-depleted RPE1 cells. Overall, this study reiterates the importance of VDAC1 in modulating tumorigenesis, due to its novel role in regulating PC length and disassembly.

## Introduction

Non-motile primary cilium (PC) is microtubule-based, membrane-ensheathed projection from cell surface, and is assembled on basal body that is transformed mother centriole[1]. PCs perform sensory functions and transduce physiological signals during development and in adult tissues[2]. Sonic Hedgehog (SHH) signalling that regulates embryonic development, tissue patterning and homeostasis in mammals is almost exclusively transduced by PC[3]. PCs are assembled in quiescent or interphase cells, and must disassemble before those cells enter mitosis. Dysfunction of PC leads to diverse multisystemic diseases that fall under ‘ciliopathies’, which also include diseases due to motile cilia dysfunction[4]. While ciliopathies are commonly associated with failed or disrupted PC assembly, ciliary disassembly defects remain largely unknown except few recent studies[5, 6]. Since PC disassembly is necessary for a cell to proliferate, sudden activation of PC disassembly in non-cycling cells may aberrantly promote cell cycle entry and support proliferation of those cells, which may ultimately facilitate tumorigenesis[7]. Importantly, loss of PC is common in epithelial cancer tissues or cancer-derived cells, though a causal relationship of PC loss and tumorigenesis is debatable and perhaps a context-dependent matter[7].

Members of Voltage Dependent Anion Chanel (VDAC) proteins are best known to localize at the outer mitochondrial membrane (OMM) and to regulate cellular bioenergetics by controlling the transport of nucleotides, small metabolites, Ca^2+^ etc between cytosol and mitochondria [8]. Recently, we identified novel centrosomal roles of all human VDACs, namely VDAC1, VDAC2 and VDAC3[9–11]. In particular, depletion of VDAC1 in growing human hTERT-RPE1 (or RPE1) led to dramatic increase in cells with PC compared to roughly 15% PC-containing control cells. Moreover, ectopic expression of GFP-VDAC1 in RPE1 cells reduced ciliation during serum starvation that is permissive condition for PC assembly[10]. Our data suggested that VDAC1 suppressed PC assembly similar to VDAC3, but mostly via non-overlapping mechanisms[10]. VDAC3 also promotes PC disassembly via cooperating with Mps1 kinase[9], which is an additional regulatory module for negative regulation of ciliogenesis.

VDAC1 is the most studied VDAC in mammals for its critical role in metabolism, interaction with hexokinase, Ca^2+^ homeostasis and regulating apoptosis[8, 12], and is considered as an attractive target for cancer therapy due to its pro-survival properties[13]. Depletion of VDAC1 in various tumor-derived cells significantly attenuated tumorigenic potential, both in cell culture and in xenograft models[14, 15]. Recent studies revealed that VDAC1 undergoes a C-terminal truncation during hypoxia likely via Legumain endopeptidase, which facilitates cellular survival via reprograming metabolism in hypoxic microenvironment[16, 17]. This form was also seen to be associated with reduced ciliation in both patient samples of clear cell renal cell carcinoma (ccRCC) or in mouse model, echoing the negative regulation of ciliogenesis by VDAC1[18, 19]. We wondered if that negative regulation of ciliogenesis by VDAC1 may be exploited against tumorigenesis of those tissues which are ciliated in healthy condition but ciliation is significantly reduced in cancer. Accordingly, we investigated here if depleting VDAC1 in various cancer-derived cells may restore ciliogenesis, which may in turn attenuate tumorigenesis in those cells.

## Materials and methods

### Cell culture

RPE1 (ATCC), Panc1 (ATCC), U87-MG (U-87; ATCC) cells were cultured in DMEM with 10% FBS (Himedia), 1% Penicillin-Streptomycin (Thermo) at 37°C in 5% CO2. For serum starvation, cells were incubated in serum free medium for indicated time. To identify cells in S-phase, cells were incubated with 50 μM BrdU for 4 h. For assessing mitochondrial health, cells were incubated with 100 nM MitotrackerRed (Invitrogen) for 45 min and fixed with 4% Paraformaldehyde (EMS). RPE1 cells were incubated with 5 μM Itraconazole (Selleckchem) or 50 μM 4,4’-diisothiocyanatostilbene-2,2’-disulfonate (DIDS; Sigma) dissolved in DMSO for indicated time. For colony formation, 2500 siRNA treated Panc1 cells were inoculated in 60 mm tissue culture dishes and were kept for 14-15 days. Cell viability was measured using XTT assay kit (Cell Signaling Technology) using a 96-well format.

### siRNA Transfection

Validated silencer select siRNAs directed against VDAC1 ORF [siVDAC1#1 (nucleotides 736-754;[10] and siVDAC1#2 (nucleotides 225-243), Ambion] and silencer negative control (Ambion) were used at 20 nM, while siRNA against TTBK2 (nucleotides 255-273; Ambion) was used at 50 nM for transfection into cells using Lipofectamine RNAiMAX (Invitrogen) following manufacturer’s instruction.

### Cytology

Cells were fixed in either 4% paraformaldehyde −0.2% TritonX-100 for 10 mins at room temperature or chilled methanol at −20° for 10 mins. Antibodies were-rabbit anti-Cep135 (1:500, Abcam); rabbit anti-Arl13B (1:250, Proteintech); mouse anti-acetylated tubulin (Ac-tub; 1:1000, Sigma); rat anti-BrdU (1:200, Abcam), rabbit anti-cleaved Caspase 3 (1:25, Abcam). Secondary antibodies were-donkey anti-rabbit or donkey anti-mouse tagged with either Alexaflour488 or Alexafluor568 (1:1000; Invitrogen) or goat anti-rat Alexaflour350 (1:200, Invitrogen). Staining for BrdU+ cells were performed as described in [11]. Images of stained cells were acquired using either a Nikon fluorescence microscope fitted with a 60X or a Zeiss Fluorescence microscope fitted with a 63X Plan Apo oil immersion objective (both NA 1.4). BrdU+ cells were imaged using a 20X objective. Some images were acquired using Leica Confocal laser scanning microscope fitted with a 63X Plan Apo oil immersion objective (NA 1.4). To determine PC length, straight or bend line was drawn from the top of the basal body to the tip of Ac-tub and Arl13B positive PC in orthogonally projected image of a ciliated cell, and the length of that line (in μ) was determined by the associated software packages (Fig.S2A; Zen blue for Zeiss or NIS-Element for Nikon).

### Immunoblotting

BCA protein assay method (Pierce) was used to determine total protein concentration of cell lysates prepared in a buffer containing 50 mM Tris-Cl pH 8.0, 150 mM NaCl and 1% NP-40 supplemented with protease inhibitors (Sigma). Antibodies for immunoblotting were-rabbit anti-VDAC (1:1000; CST), mouse anti-α-Tubulin DM1A (α-tub; 1:10,000; Sigma), rabbit anti phospho-AMPK (1:1000; CST), mouse β-actin-conjugated with DyLight800 (1:10000; Biorad). Alexafluor680-conjugated donkey anti-rabbit and DyLight800-conjugated donkey anti-mouse (both 1:10000; Invitrogen) secondary antibodies were used for fluorescence scanning using Odyssey-CLX infrared imaging system (Li-Cor). For AurkA, HRP-conjugated goat anti-rabbit secondary antibody was used, followed by chemiluminescence-based imaging using GelDoc MP system (Biorad). The background-corrected intensities of bands were determined using ImageStudio (Li-cor).

### qRT-PCR

Total RNA from cells was isolated using TRIzol (Invitrogen) according to the manufacturer’s protocol, and dissolved in 10 μL of nuclease free water. 1 μg DNase-treated total RNA was then reverse transcribed using iScript cDNA synthesis kit (Biorad). The cDNA was subsequently used for quantitative, real-time PCR (qPCR) using SsoAdvanced Universal SYBR Green Supermix (BioRad) on a CFX Connect Real Time system (BioRad) where the mentioned oligos were used at a concentration of 500 nM each. qPCR primer-pair efficiency (E) of all sets of primers (GAPDH_Fwd: GTCTCCTCTGACTTCAACAGCG, GAPDH_Rev: ACCACCCTGTTGCTGTAGCCAA; VDAC1_Fwd: CTCAGCCAACACTGAGACCA, VDAC1_Rev: TGCAAGCTGATCTTCCACAG; VDAC2_Fwd: CAGTGCCAAATCAAAGCTGA, VDAC2_Rev: CCTGATGTCCAAGCAAGGTT; VDAC3_Fwd: CGTTGGTTTGAAGACCTTCAGCGT, VDAC3_Rev: GGCTCTTATGAAACCAAAGACCTCCACC) were calculated using manufacturer’s protocol (Biorad; E-values ranging from 90%-110% were considered as efficient primer pairs).

## Results and Discussion

### Depletion of VDAC1 increases PC frequency and length of those PCs in cancer-derived cells

PCs are rare or absent in most solid tumor tissues and consequently in tumor-derived epithelial cells, though a causal role of absence of PC and tumorigenesis is not established in some cancers[7]. Previous studies indicated that epithelial PC were gradually shortened and lost from various types of pancreatic cells during the tumorigenic progression leading to pancreatic ductal adenocarcinoma (PDAC), one of the most lethal cancers[20, 21]. VDAC1 is overexpressed in various cancer tissues including pancreatic cancer[22, 23]. We show higher mRNA expression of VDAC1 in Panc1, a PDAC-derived cell, as compared to RPE1, a non-cancerous diploid cell (Fig. S1A). Since VDAC1 suppresses PC assembly in growing RPE1 cells[10], and pancreatic epithelial tissues lost PC during tumorigenesis [21, 24], we sought to examine if depleting VDAC1 may increase ciliation in Panc1 cells.

We depleted VDAC1 in Panc1 cells using the two siRNAs, the one that was previously used (siVDAC1#1) in[10], and a second siRNA (siVDAC1#2), to validate that ciliary phenotypes observed in these two treated cells are indeed due to VDAC1 depletion. While both siRNAs effectively downregulated VDAC1 (80-90% mRNA downregulation; Fig.S1A and 65-70% protein depletion; Fig.1B, S1B), siVDAC1#1 appeared as more potent. Neither of these siRNAs affected the expression of VDAC2 and VDAC3, validating the specific binding of these siRNAs to VDAC1 mRNA (Fig.S1A). Though VDACs are critical for bioenergetics, none of the VDACs is essential for cellular respiration or mitochondria-mediated death[25, 26], likely due to functional compensation by other VDACs. Expectedly, VDAC1 depletion in Panc1 cells did not affect overall mitochondrial health as judged by mitotrackerRed incorporation in siVDAC1 cells and siControl cells (Fig.S1C).

**Fig 1.**
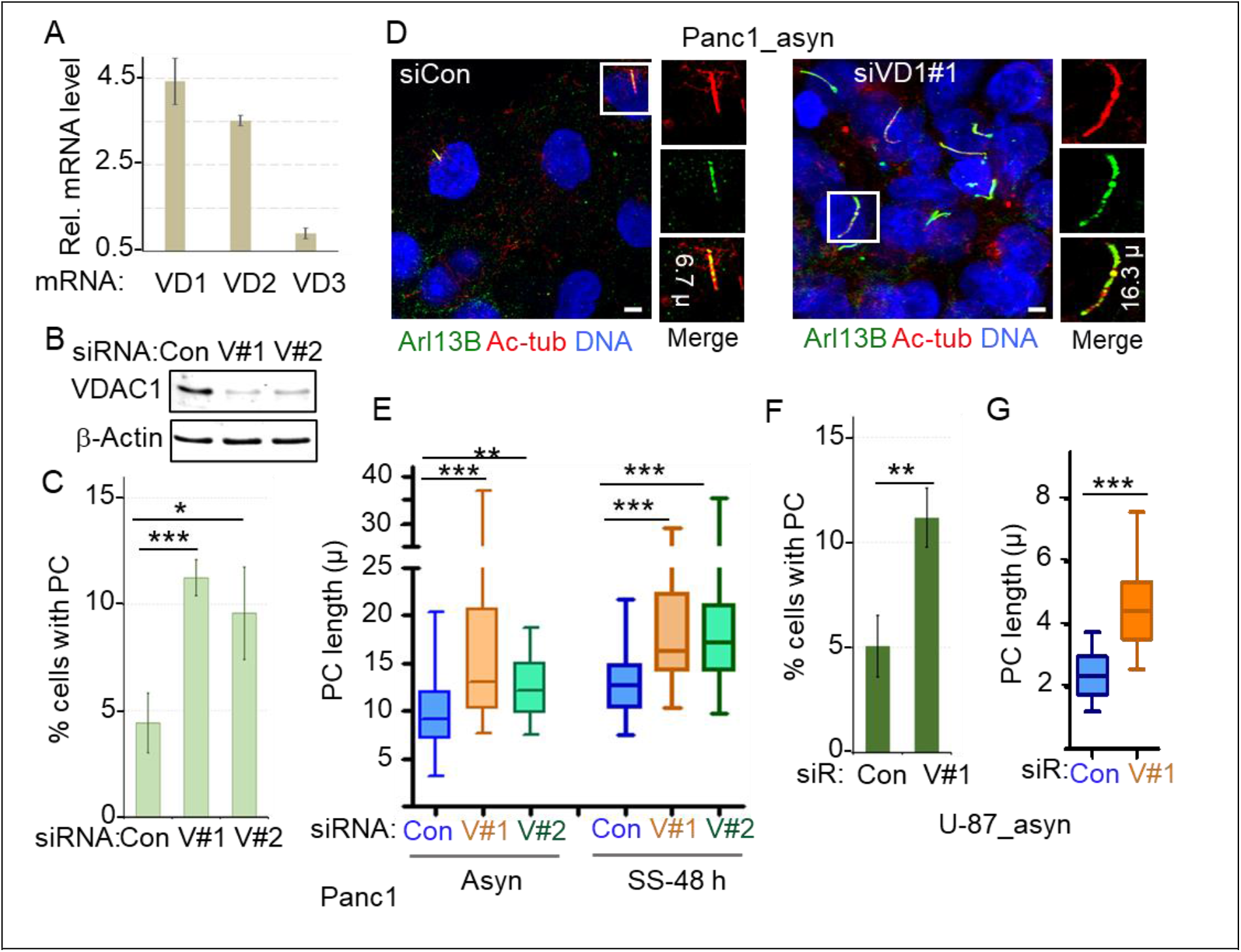
VDAC1 depletion leads to increased ciliation and drastic elongation of PC in cancer-derived cells. (A) The ratio of GAPDH normalized mRNA levels of human VDAC1 (VD1), VDAC2 (VD2) and VDAC3 (VD3) genes in Panc1 cells to that in RPE1 cells were measured by qRT PCR, and are plotted as bars, n= 3. (B-D) Asynchronously growing Panc1 cells were treated with VDAC1-specific siRNAs (siVDAC1#1 or V#1 or siVDAC1#2 or V#2) or control siRNA (Con) for 72 h. (B) Representative immunoblots show depletion of VDAC1 in siRNA-treated Panc1 cells, where β-actin was used as loading control. (C) PCs in Panc1 cells were identified by staining for acetylated tubulin (Ac-tub) and Arl13B. Percentage of cells with PC were plotted as bars where values represent mean ± S.D. from three independent experiments for which 350–400 cells were counted per replicate. Here and in all other cases, p value was determined by unpaired t-test, where *** indicates p<0.001, ** indicates p < 0.01, * indicates p < 0.05 and n.s. indicates non-significant. (D) Representative micrographs show random fields of siCon and siVD1#1 cells. DNA is blue, bar= 5 μ. Insets show two-fold magnified PC, and the length of that PC is shown. (E) The length of PC (in μ) of 40 randomly picked Panc1 cells treated with siRNAs and incubated in serum-containing or serum-starved medium were determined, and are presented in a box and whisker diagram. Here and in other experiments, boxes indicate lower and upper quartiles, the marker in the box (–) indicates the median, and the whiskers represent minimum and maximum values for each series. (F-G) Asynchronously growing U-87 cells were treated with siVDAC1#1 (V#1) or siCon (Con) for 72 h, and PCs were identified by staining for markers of cilia and basal bodies. (F) Percentage of cells with PC were plotted as bars where values represent mean ± S.D. from three independent experiments for which 450–500 cells were counted per replicate. (G) The length of PC (in μ) of 40-45 randomly picked ciliated U-87 are presented in a box and whisker plot.

Next, we examined the PCs in siRNA-transfected Panc1 cells using Arl13B and Ac-tub as the ciliary markers. Normally, 3-5% of an asynchronously growing population of Panc1 cells contain PC, while it increases to 10% when incubated in serum-starved medium for 48 h(Fig.1C-D, S2B-C; [21, 24]). Interestingly, depletion of VDAC1 increased the ciliation in Panc1 cells to roughly 10-12%in growing condition i.e. in presence of serum, and roughly 25% in serum-starved condition (Fig.1C-D, S2B-C). A similar increase in ciliation is also observed in U-87 glioblastoma-derived cells upon VDAC1 depletion (Fig.1F, S2D). We chose to test glioblastoma-derived cells as PCs were observed, albeit infrequently, in glioblastoma-derived cells [27] and in patient-derived glioblastoma tumorsphere [28]. This observed increase in ciliation upon VDAC1 depletion suggests a critical role of VDAC1 in suppressing ciliogenesis in these cancer-derived cells that assemble sporadic PC, say 2-5%. We also examined other cell lines derived from diverse solid tumors, which showed none or less than 1% PC in growing condition. However, we only observed slight increase in ciliation upon VDAC1 depletion in those cells that had less than 1% PC in control transfection (data not shown). We did not include these cells in our detailed analysis, as it would be non-significant. Notably, increased ciliation were observed in a significant yet a small fraction of Panc1 and U-87 cells upon VDAC1 depletion, raising the possibility that the observed effect may not be linked to the altered metabolic status of these VDAC1-depleted cells as proposed earlier [23, 29, 30], which would possibly show a greater readout. Also, VDAC1 depletion might not cause a chronic attenuation in cellular ATP level as judged by only a marginal increase in the conserved phosphorylation of AMP-activated protein kinase α in siVDAC1 treated Panc1 cells compared to control (Fig.S1D). A rather plausible explanation could be the loss of function of a pool of VDAC1 that localizes to the centrosomes/basal bodies as shown earlier[10].

Surprisingly, VDAC1 depletion drastically increased the length of PC in Panc1 cells, in both asynchronously growing or serum starved conditions (Fig.1E). Commonly, PCs in Panc1 cells are considerably longer than that are formed in RPE1 cells (on average 5-8 μ in Panc1 versus 2.5-4 μ in RPE1). In contrast, PCs formed in VDAC1-depleted Panc1 cells were mostly longer than 10 μ, and in some cases 25-30 μ, which is, to the best of our knowledge are not seen in any other cell types. PCs were also significantly longer in VDAC1-depleted U-87 cells compared to the control (Fig.1G). Such remarkable elongation of PC upon VDAC1 depletion in both cell types suggest that VDAC1 likely regulates molecular pathways that directly regulates PC length, and such effect is perhaps not due to aberrant metabolic status or cell cycle stage. However, delineation of the mechanistic details of how cilia length are regulated by differentially localized VDAC1 pool is not within the scope of this study.

### Depletion of VDAC1 delays cell cycle and attenuates proliferation of Panc1 cells

Commonly, PC disassembly starts during S-phase entry, and is completed before G2/M transition. The presence of PC allows regulated activation of G1-S modulators and timely progression of cell cycle, while loss of PC may result in aberrant cell cycle that supports higher proliferation and promotes other tumorigenic factors[7, 31]. Panc1 cells are proliferating cancer-derived cells with only sporadic PC. To examine if increased ciliation in VDAC1-depleted Panc1 cells would affect cell cycle, we counted the S-phase cells judged by their ability to incorporate BrdU in asynchronous population 72 h post-siRNA transfection. We observed a significant decrease in BrdU+ Panc1 cells treated with VDAC1-specific siRNAs compared to the control cells, indicating a delay in S-phase entry (Fig.2A-B). A few BrdU+ siVDAC1 cells even contained mature PC that were positive for both Ac-tub and Arl13B (not shown), while no such cell was seen in control. Since ciliation was greater in serum-starved Panc1 cells, we examined starved cells for BrdU incorporation, considering the fact that serum starvation does not restrict Panc1 cells to quiescence. Roughly 10-15% starved siControl cells incorporated BrdU. However, further decrease in BrdU incorporation in VDAC1-siRNA treated cells was not significant (Fig.S3B). Thus, VDAC1 depletion led to a delay in cell cycle progression in proliferating Panc1 cells. Such cell cycle delay due to enhanced ciliation might affect mitotic entry as well. Therefore, we examined the population of mitotic cells judged by condensed chromosomes and the presence of mitotic spindle. We observed a significant lowering in the mitotic index in siVDAC1-treated Panc1 cell population than the control cells (Fig.2E). Overall, these two phenomena likely contributed to a moderate decrease in viability of VDAC1-depleted Panc1 cells (Fig.3C). VDAC1 is implicated in mitochondria-mediated apoptosis, as its oligomerization at the mitochondrial outer membrane releases apoptotic stimuli such as Cytochrome C[8]. However, VDAC1 depletion was shown to attenuate various stimuli-induced apoptosis. Here, we did not observe any change in intrinsic apoptosis in VDAC1-depleted Panc1 cells than the control, as judged by cleaved-caspase 3 positivity, while rotenone treatment increased the percentage of apoptotic Panc1 cells similar to the published data (Fig.S2A, [32]). VDAC1-siRNA treated Panc1 cells were unable to form colonies upon limiting dilution, as compared to the control cells (Fig.2D). Reduction in cell proliferation upon VDAC1 downregulation in various cancer-derived cells were demonstrated earlier, which ultimately led to reduced cell motility and cancer cell growth, using both in vitro and in vivo xenograft studies[15, 33, 34]. Our data agrees well to those observations, and adds that a significant delay in cell cycle likely contributes to the cause of reduced cell growth and viability, and thereby attenuated colony formation abilities of VDAC1-depleted Panc1 cells.

**Fig 2.**
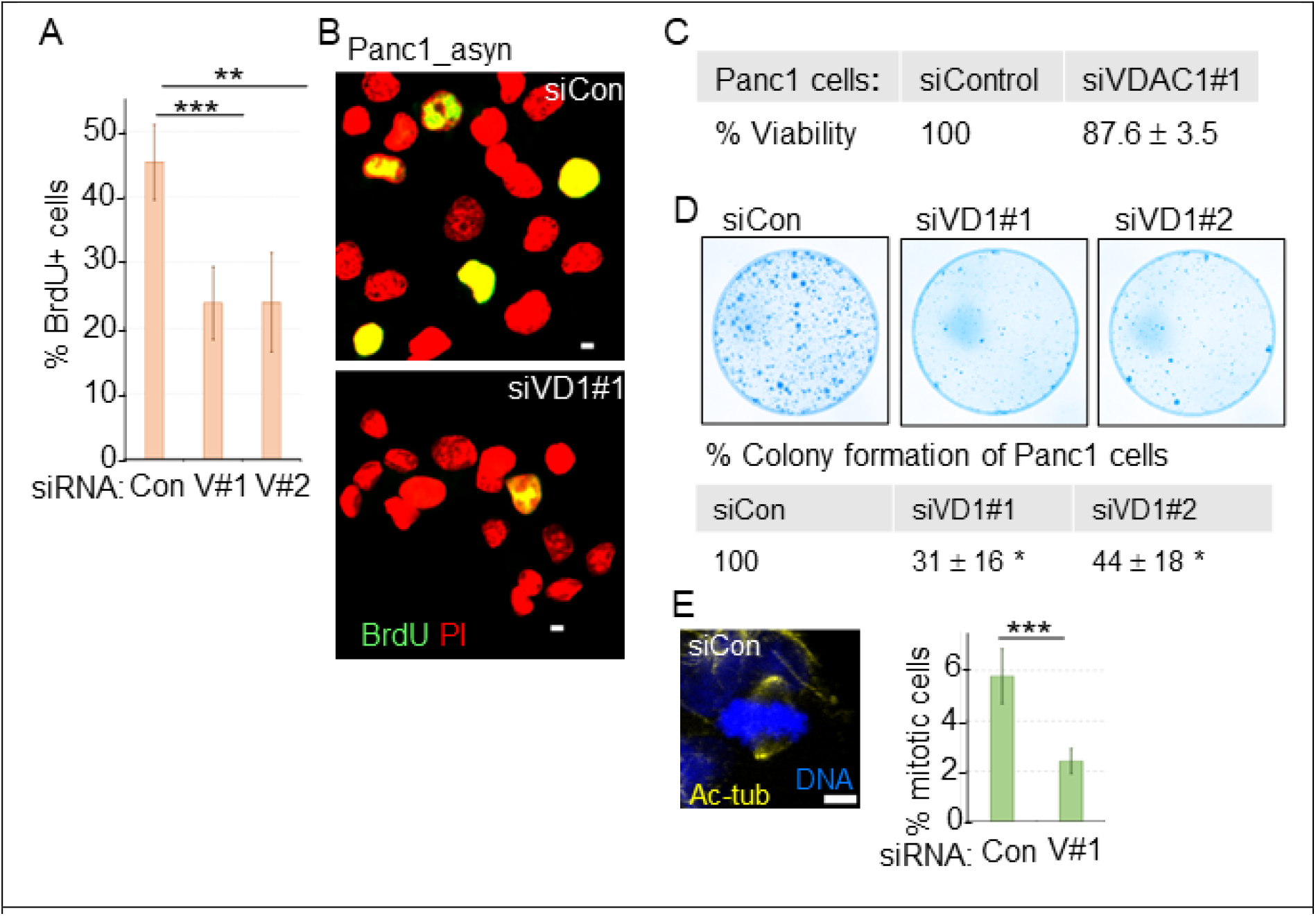
VDAC1 depletion in Panc1 cells reduced S-phase cells and inhibited proliferation. (A-B) Asynchronous population of Panc1 cells transfected with VDAC1-specific siRNAs (V#1 and V#2) and control siRNA (Con) were grown for 68 h, and then 4 h in presence of BrdU. (A) The percentage of BrdU+ cells were plotted as bars where values represent mean ± S.D., n=3; 300-400 cells were counted per replicate. (B) Micrographs show random fields of siControl (siCon) and siVDAC1#1 (siVD1#1) cells stained for BrdU (green), where DNA is stained by propidium iodide (PI, red). Bar= 10 μ. (C) Asynchronously growing Panc1 cells treated with control and VDAC1#1 siRNAs for 72 h were examined for cell viability using XTT assay. % Viability of indicated cells are shown in the table where values represent mean ± S.D., n=3. (D) Representative images of crystal violet-stained colonies formed by Panc1 cells treated with indicated siRNAs and inoculated in a limiting dilution for two weeks. The table shows the colony counting analyses, where values represent mean ± S.D., n=3. (E) siRNA transfected Panc1 cells were examined for the presence of mitotic cells judged by condensed chromosomes and Ac-tub-stained spindles. Percentage of mitotic cells are shown as the bars where values represent mean ± S.D., n=3 and at least 400 cells were analyzed per replicate. A representative micrograph of such mitotic cell is shown, bar = 5 μ.

**Fig 3.**
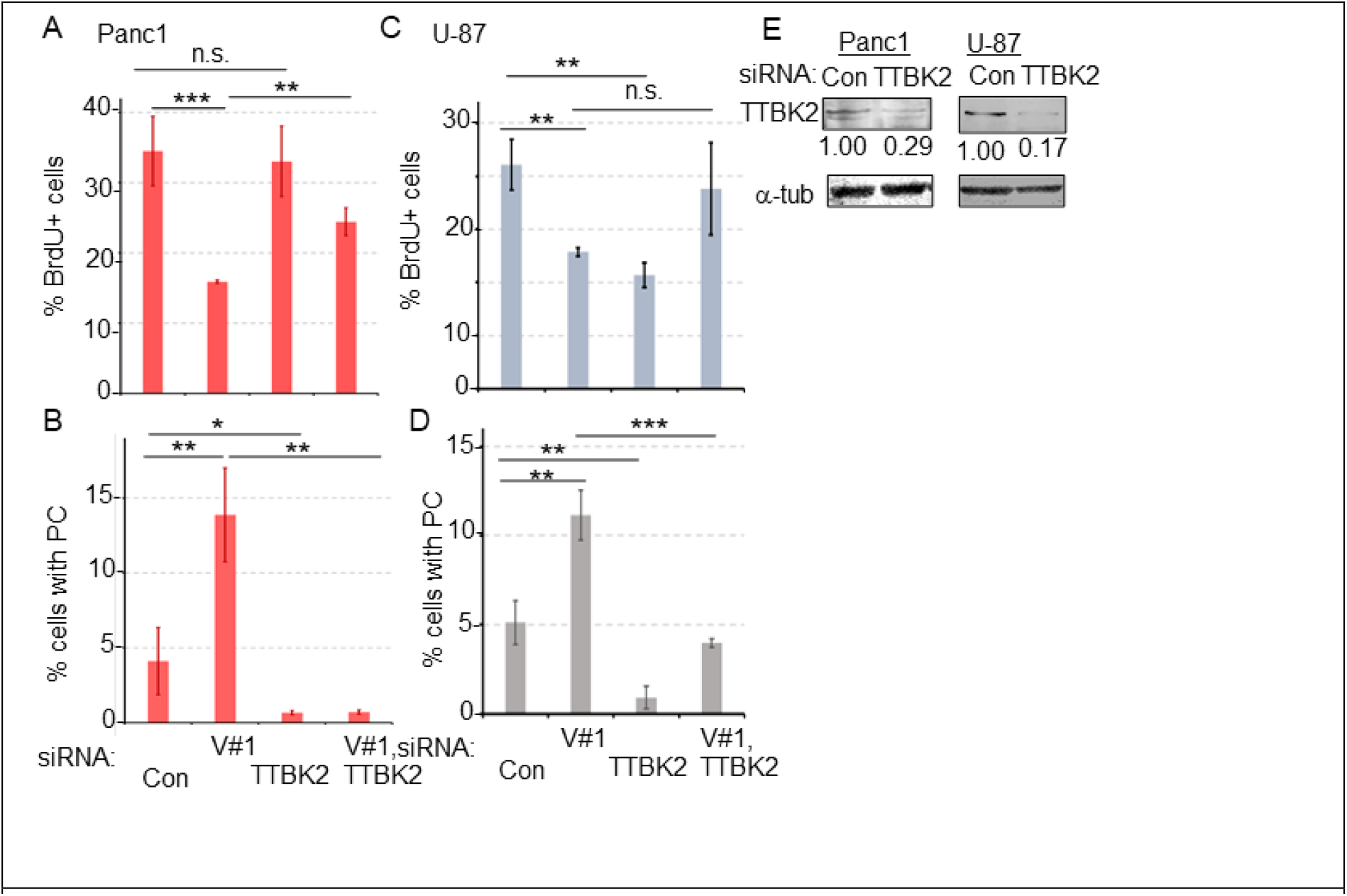
Inhibiting PC formation restored cell cycle delay in VDAC1-depleted cancer-derived cells. (A, C) BrdU incorporation was examined in growing Panc1 cells (A) or in U-87 cells (C) transfected with indicated siRNAs (alone or in combination). The percentage of BrdU+ cells are plotted as bars where values represent mean ± S.D., n=3, at least 500-700 cells were counted per replicate. (B, D) Similarly treated cells were examined for the presence of PC marked by Arl13B and Ac-tub in Panc1 cells (B) or in U-87 cells (D), and the percentage of PC are plotted as bars where values represent mean ± S.D., n=3, at least 400 cells were examined per replicate). (E) Efficiency of TTBK2 depletion by siTTBK2 in Panc1 and U-87 cells were determined by immunoblotting. Representative immunoblots are shown, where values indicate normalized intensities of TTBK2 bands, where μ-tub was used as loading control.

### Inhibition of PC assembly in VDAC1-depleted cells restored the delay in S-phase entry

Our data indicated that the increase in cancer cells with PC were likely due to elimination of PC assembly suppression ability of VDAC1, which poses a delay in S-phase entry in these cells. If so, then abolishing the PCs in VDAC1-depleted cells should restore cell proliferation. Therefore, we sought to inhibit the pathway required for core PC assembly by targeting Tau tubulin kinase 2 (TTBK2) that is required for PC biogenesis[35, 36]. TTBK2 is located to the distal appendages and helps the removal of CP110 from the axoneme tip during PC assembly initiation[37]. Accordingly, TTBK2 depletion abolishes PC even in permissive condition for PC assembly[38]. We tested a validated siRNA against TTBK2[38] in serum-starved RPE1 cells, where, a modest depletion of TTBK2 siRNA led to remarkable decrease in ciliation (Fig. S3A-B). Next, we treated asynchronously growing Panc1 and U-87 cells with siVDAC1#1 alone, and in combination with siTTBK2. Expectedly, VDAC1 depletion increased ciliation, while TTBK2 depletion drastically reduced ciliation in both Panc1 and U-87 cells (Fig. 3B, D-E). When Panc1 cells were treated with both siRNAs, less than 1% cells had PC suggesting that cells could not assemble PC due to loss of TTBK2 function (Fig. 3B). Importantly, BrdU incorporation was significantly restored in those double siRNA transfected Panc1 cells compared to that in VDAC1-depleted cells (Fig. 3A), suggesting that loss of PC in those cells likely releases the cell cycle delay leading to increase in S-phase cells. We observed a similar trend in BrdU positivity in U-87 cells as well, where it modestly increased in cells treated with both siRNAs compared to cells treated with siVDAC1#1 alone. However, the changes in ciliation and proliferation in U-87 cells were not as prominent as in Panc1 cells, with low margin of error. Moreover, regulation of TTBK2 expression and function in glioblastoma cells is complex due to the presence of circular RNA (circ-TTBK2) generated from the first few exons of TTBK2 gene, and the siRNA used here might have modulated that circ-TTBK2[39]. Such regulation may ultimately decrease the proliferation of U-87 cells[39], which might have impacted the readout of our experiment. Nevertheless, our data in these two cell types indicated that if PC assembly is attenuated in VDAC1-depleted cells, cell cycle delay can be significantly restored. Thus, we demonstrated that downregulation of VDAC1 in a population of cancer-derived cells that assembles infrequent PC, increases ciliation leading to delayed cell cycle, which contributes to reduced proliferation, a prominent tumorigenic potential of these cells. Our study complements well with the earlier studies of VDAC1-depletion in PDACs and glioblastoma, where altered metabolism due to VDAC1-depletion was considered as the driving factor of attenuating tumorigenic potential[15]. Interestingly, VDAC1-depletion in glioblastoma was considered to modulate various signaling pathways and to increase differentiation potential[30, 33]. It is possible that VDAC1-depleted GBM-derived cells might assemble PC, which facilitated differentiation. Overall, It is tempting to speculate that overexpression of VDAC1 in these cancer cells contributed to proliferation and other tumorigenic properties, at least partly by suppressing the PC assembly.

### Targeting VDAC1 inhibits serum-induced PC disassembly and increases PC length in RPE1 cells

Depletion of VDAC1 allowed a small, yet significant population of some cancer-derived cells to assemble PC in growing condition. A similar, but higher increase in ciliation was observed in asynchronously growing RPE1 cells upon VDAC1 depletion[10]. Moreover, overexpression of GFP-VDAC1 in RPE1 cells during serum starvation, the permissive condition of PC assembly, significantly suppressed ciliation[10]. Here, we additionally discovered VDAC1 to regulate PC length in cancer-derived cells. Since the magnitude of ciliation in these cells is small and since these cancer-derived cell populations are heterogenous, we sought to examine how VDAC1 regulates PC length in RPE1 cells, which synchronously enter cellular quiescence, and disassemble PC synchronously upon serum addition, using a standardized strategy (Fig.4A).

**Figure 4.**
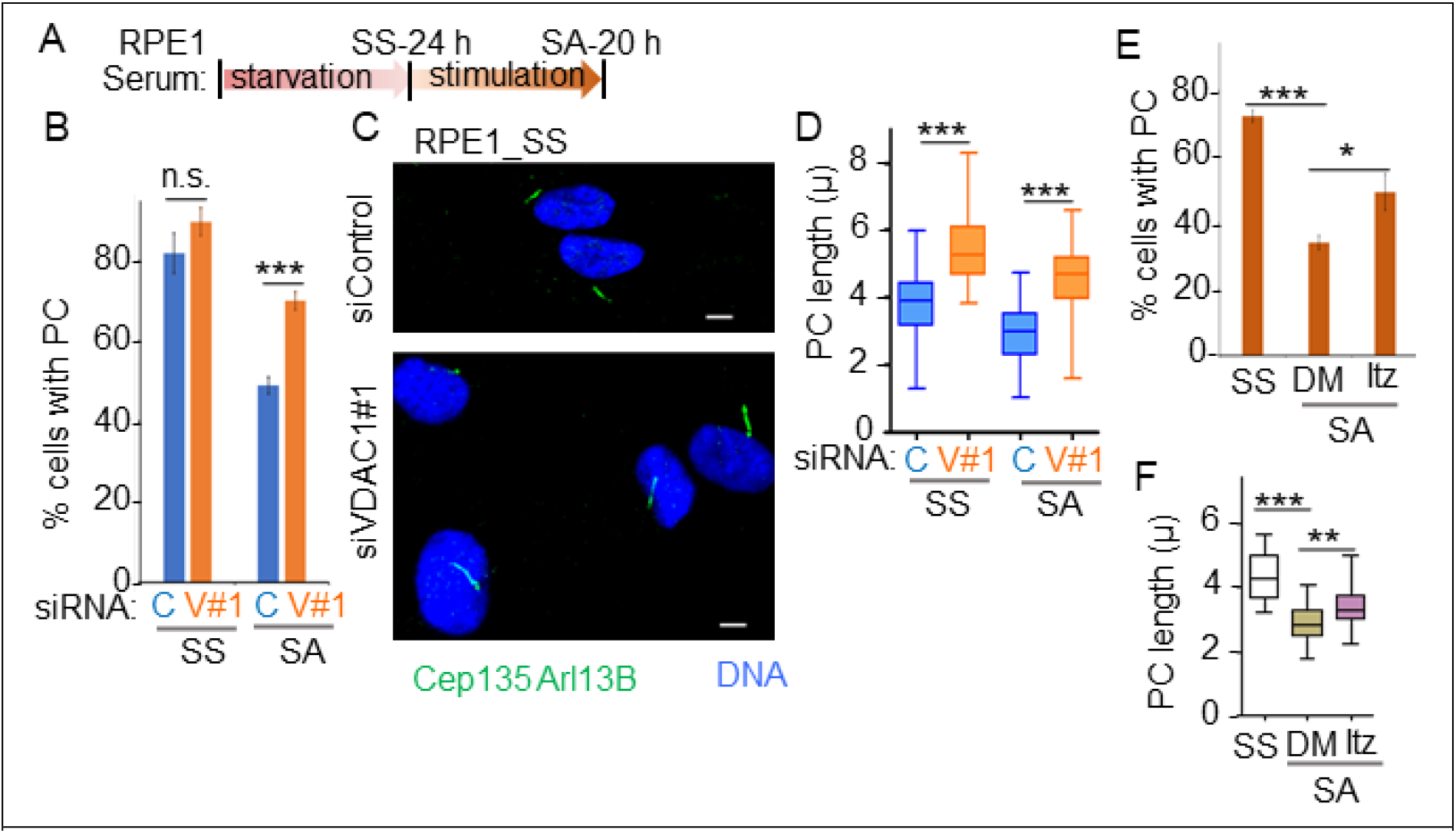
Targeting VDAC1 increases PC length in RPE1 cells during quiescence and also during serum-induced PC disassembly in RPE1 cells. (A) Cartoon of the experimental strategy to serum-starve RPE1 cells for 24 h (SS) followed by serum addition and examining those cells after 20 h (SA). (B-D) RPE1 cells treated with siCon or siVDAC1#1 were examined for ciliation after serum starvation (SS) and during serum addition (SA), and the percentage of cells with PC were plotted as bars (B) where values represent mean ± S.D., n=3; 250–300 cells were counted per replicate. (C) Representative micrographs show random fields of indicated cells stained for PC and basal body (both are green). DNA is blue, bar= 5 μ. (D) The length of PC (in μ) of 40 randomly picked siRNA-treated RPE1 cells are presented in a box and whisker diagram (E-F) RPE1 cells were treated with either Itraconazole (Itz) or DMSO (solvent control) during SA and stained for PC. (E) Percentage of cells with PC are plotted as bars where values represent mean ± S.D., n=3, with at least 300 cells were counted per replicate. (F) The length of PC (in μ) of 40-50 random, serum starved, and serum-added with Itz or DMSO treated RPE1 cells were determined, and are presented in a box and whisker plot.

Accordingly, we depleted VDAC1 in serum-starved RPE1 cells. PC was observed in roughly 80-85% of serum-starved siControl and siVDAC1#1 cells (Fig. 4B). The extent of ciliation could not alter much upon VDAC1 depletion in serum-starved condition. Importantly, PCs were significantly longer in VDAC1-depleted RPE1 cells compared to the control cells (Fig. 4C-D), corroborating our observation in cancer-derived cells. Serum-induced PC disassembly takes place in a regulated manner that involves shortening of PC. We previously showed that after 20 h of serum re-addition, roughly 50% of the control RPE1 cells resorbed their PC[9]. If VDAC1 regulates PC length then VDAC1 depletion would prevent that shortening of PC during serum-induced PC disassembly in RPE1 cells. We observed a significant increase of VDAC1-depleted cells to retain PC after serum addition compared to the control cells suggesting a novel role of VDAC1 to promote PC disassembly (Fig. 4B). Importantly, majority of the PCs in siVDAC1#1-treated RPE1 cells were longer than the control cells during that condition (Fig. 4D). Since VDAC1-depleted cells had longer PC during starved condition, possibly the average length of the retained PCs were longer in those cells even after serum stimulation.

Itraconazole (Itz) inhibits angiogenesis via binding to VDAC1 and likely inhibiting its channel activity and ultimate decreasing cellular ATP level[40]. That results in acute increase in AMPK activity, which attenuates mTOR activity and cellular proliferation in endothelial cells[40]. Separate studies showed that itraconazole inhibits Hedgehog signalling by binding to smoothened and inhibiting its localization to PC[41]. We wondered if itraconazole may inhibit VDAC1 during serum-induced PC disassembly, and to affect PC length. Accordingly, RPE1 cells were treated with itraconazole during the onset of serum addition, and ciliation were examined after 20 h. Itraconazole treatment prevented PC disassembly and also led to increase in PC length, however quite modestly (Fig. 4E-F), as compared to VDAC1 downregulation, suggesting that itraconazole-mediated chronic inhibition of VDAC1’s channel activity had minimal influence on ciliogenesis. Also, such long-term treatment of these cells with itraconazole did not significantly alter the cellular ATP balance (Fig. S4D). Possibly, the two other VDACs compensated the cellular ATP level during inhibition of VDAC1’s channel activity. We also tested DIDS, another inhibitor of VDAC1, which particularly inhibits the oligomerization of VDAC1 induced by apoptotic stimuli, in regulating PC length. Since RPE1 does not overexpress VDAC1, treating cells with DIDS did not affect ciliation or PC length during serum-induced PC disassembly (Fig. S4E-F). Overall, our data suggest that VDAC1 plays a role in PC disassembly, and more critically in regulating PC length.

VDAC1 is the best studied for its role in cellular bioenergetics and its high expression is associated with low-survival rate in many cancers suggesting it as an attractive prognostic marker and therapeutic target [14, 29, 30, 42–44]. We first identified VDAC1 to localize to centrosomes/basal bodies, and to regulate ciliogenesis[10]. Here, we proved that VDAC1 promotes PC disassembly and is critical in regulating PC length, in both normal and cancer-derived cells establishing VDAC1 as an important regulator of ciliogenesis. Since, mitochondrial dysfunction or balance between mitochondrial fission-fusion may affect ciliation[45, 46], it may not be surprising that VDAC1-depletion may regulate ciliogenesis. However, presence of aberrantly long cilia in VDAC1-depleted cells is surprising. It can be speculated that aberrant PC elongation is likely due to the loss of VDAC1’s interaction with the molecular players of ciliogenesis that directly regulate PC length. Though elucidating the mechanism of how VDAC1 may regulate PC length is beyond the scope of this study, it may be noted that dynein-light chain Tctex1 that regulates PC length via a dynein-independent mechanism is an interactor of VDAC1[47, 48]. Our future studies will shed light on this molecular interaction in regulating PC length, and how targeting VDAC1-regulated ciliogenesis may ameliorate therapeutic strategies against tumorigenesis.

## Acknowledgement

This work is supported by Ramalingaswami fellowship to SM by the Department of Biotechnology (DBT), Govt. of India, research grant from Department of Science & Technology and Biotechnology, Govt. of West Bengal (525(Sanc.)/ST/P/S&T/2G-4/2018) to SM, the core research grants from the Science and Engineering Research Board (SERB), Govt. of India separately to SM (CRG/2020/004042) and CM (CRG/2019/006427). AD and PH are respectively DBT and UGC research fellows. Authors sincerely thank Drs Virupakshi Soppina, IIT Gandhinagar; Piyali Mukherjee, Presidency University; Oishee Chakrabarti, Saha Institute of Nuclear Physics for reagents, and Abhik Saha, Presidency University for providing Odyssey-CLX facility. Authors are thankful to Dr Pralay Majumder and Priyanka Das for technical help, and the microscope facility of Presidency University.

## Figures and Legends

**Fig. S1.**
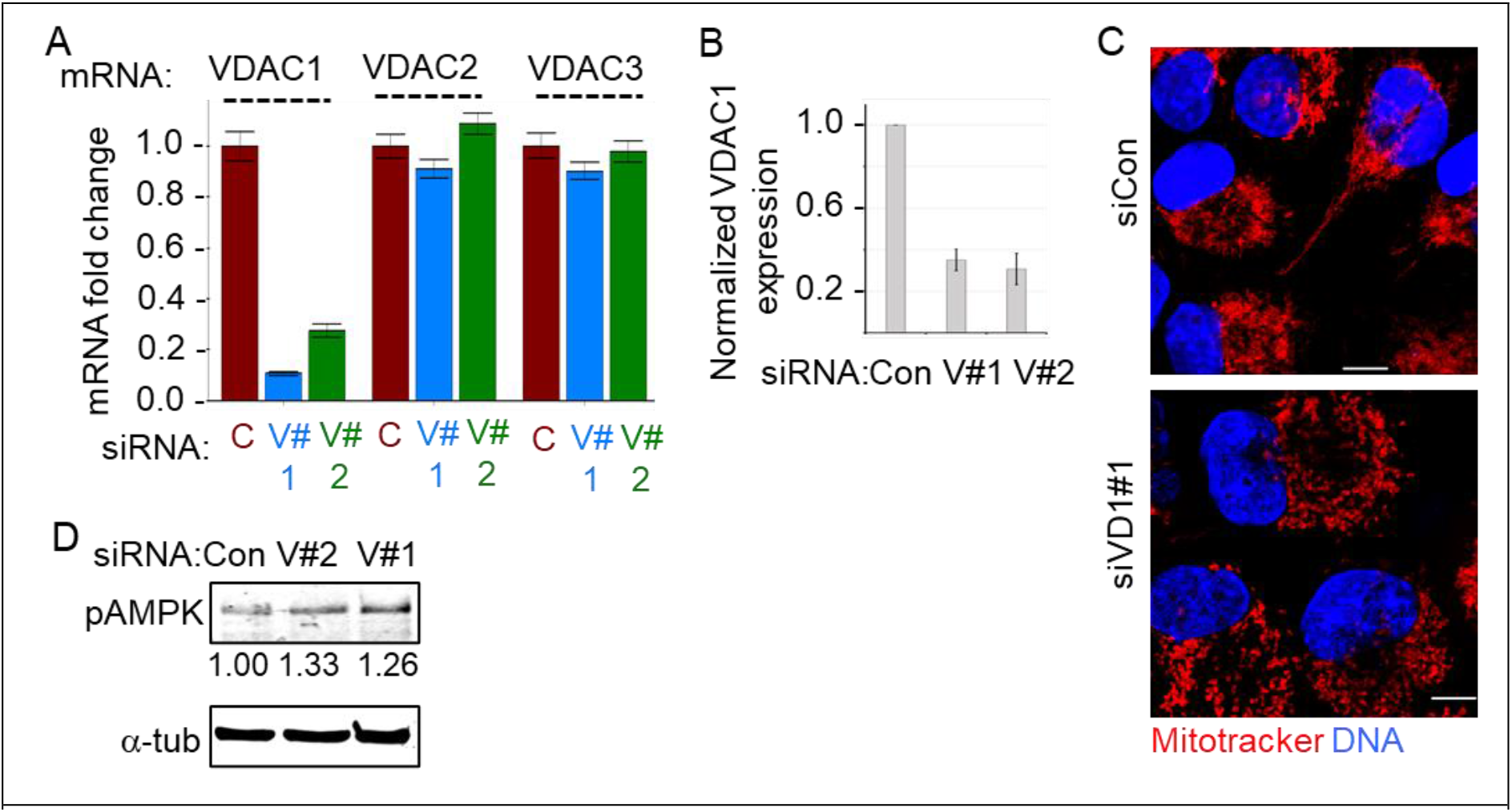
Asynchronously growing Panc1 cells were treated with control (C) and VDAC1-specific (V#1 and V#2) siRNAs for 72 h. (A) Relative mRNA levels (fold change) of three human VDACs were measured by qRT-PCR using GAPDH as normalizing control, from those cells and plotted as bars, n=3. (B) Depletion of VDAC1 protein from that experiment was determined using immunoblots (a representative blot is shown in Fig.1B), using β-actin as loading control. Bars show the normalized intensities of VDAC1 band, where values represent mean ± S.D. (n=3). (C) Representative confocal micrographs show mitotracker red incorporation in Panc1 cells treated with indicated siRNAs. DNA is blue, bar is 10 μ. (D) Representative immunoblots show the level of phospho-AMPK (pAMPK) in Panc1 cells treated with indicated siRNAs for 72 h. Values show normalized intensities of phospho-AMPK, where α-tub was used as loading control.

**Fig. S2.**
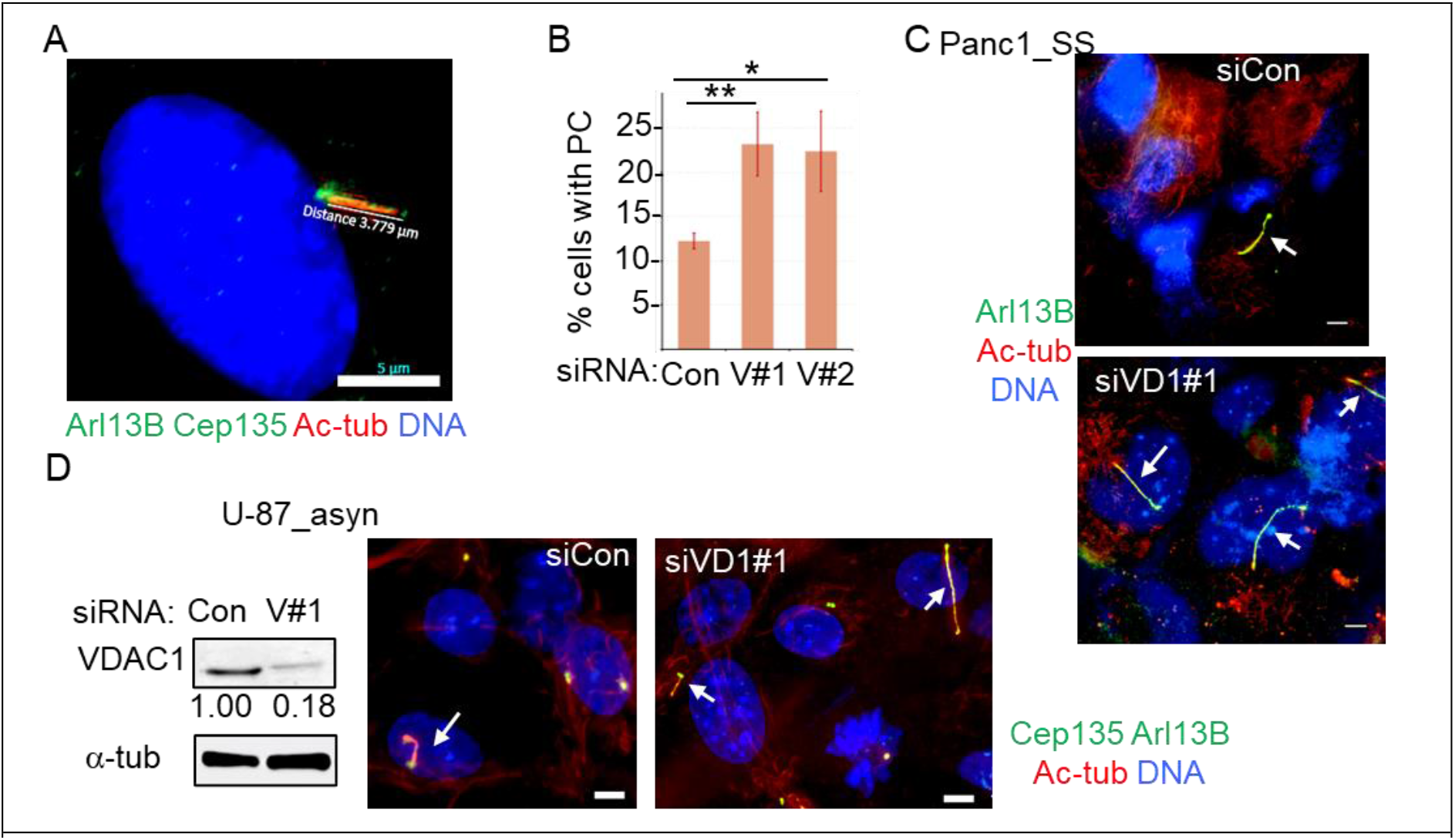
(A) Representative image of a serum starved RPE1 cell stained with antibodies against basal body and ciliary proteins, where the length of PC was measured (length is 3.779 μ) by drawing a line from the base (top of the basal body) to the tip of the PC using the Zen Blue software associated with Zeiss fluorescence microscope. DNA is blue, bar =5 μ. Similar technique was used to measure PC length throughout the study in RPE1, Panc1 and U-87 cells. (B) Panc1 cells treated with indicated siRNAs were incubated in serum-starved medium for 48 h, and the PC were analysed similarly as in Fig.1D. Percentage of cells containing PC are plotted as bars where values represent mean ± S.D., n=3, 350–400 cells were counted per replicate. (C) Representative micrographs from that experiment show random fields of Panc1 cells stained for PC (indicated by arrow) markers. DNA is blue, bar = 5 μ. (D) Asynchronously growing U-87 cells were treated with control (siCon) and VDAC1-specific (siVD1#1) siRNAs for 72 h. Representative immunoblot shows the depletion of VDAC1, where values show normalized intensities of VDAC1 band using α-tub as loading control. (E) Representative micrographs from that experiment show random fields of U-87cells stained for the indicated antibodies, where DNA is blue, bar = 5 μ. Arrow indicates PC.

**Fig. S3.**
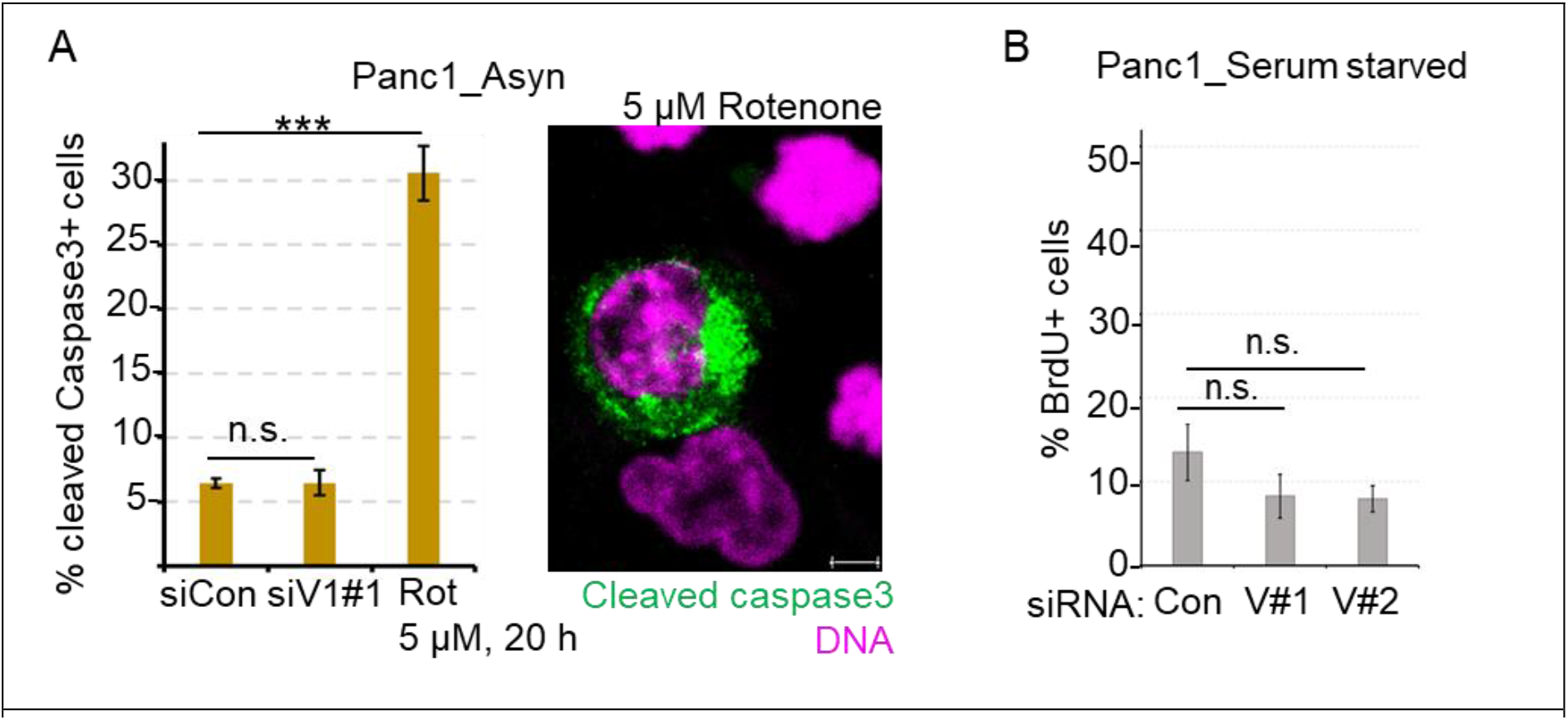
(A) Asynchronously growing Panc1 cells were treated with 5 μM rotenone for 20 h following the published protocol in Sarkar et al., 2022; FEBS Journal. Cells were then fixed and stained using specific antibody against cleaved Caspase3 as the marker of apoptotic cell. Representative image shows a random field of rotenone treated Panc1 cells, where only an apoptotic cell is stained green; bar is 5 μ. Percentage of cells positive of cleaved Caspase3 were analyzed and plotted as bars, where values represent mean ± S.D., n=3, 900–1100 cells were counted per replicate. Similarly growing Panc1 cells treated with control (siCon) and VDAC1-specific (siV1#1) siRNAs for 72 h, were also analyzed for apoptotic population, and are plotted similarly as bars. (B) Percentage of Brdu+ Panc1 cells treated with indicated siRNAs and serum-starved for 48 h are plotted as bars where values represent mean ± S.D., n=3; 300-400 cells were counted per replicate.

**Fig. S4.**
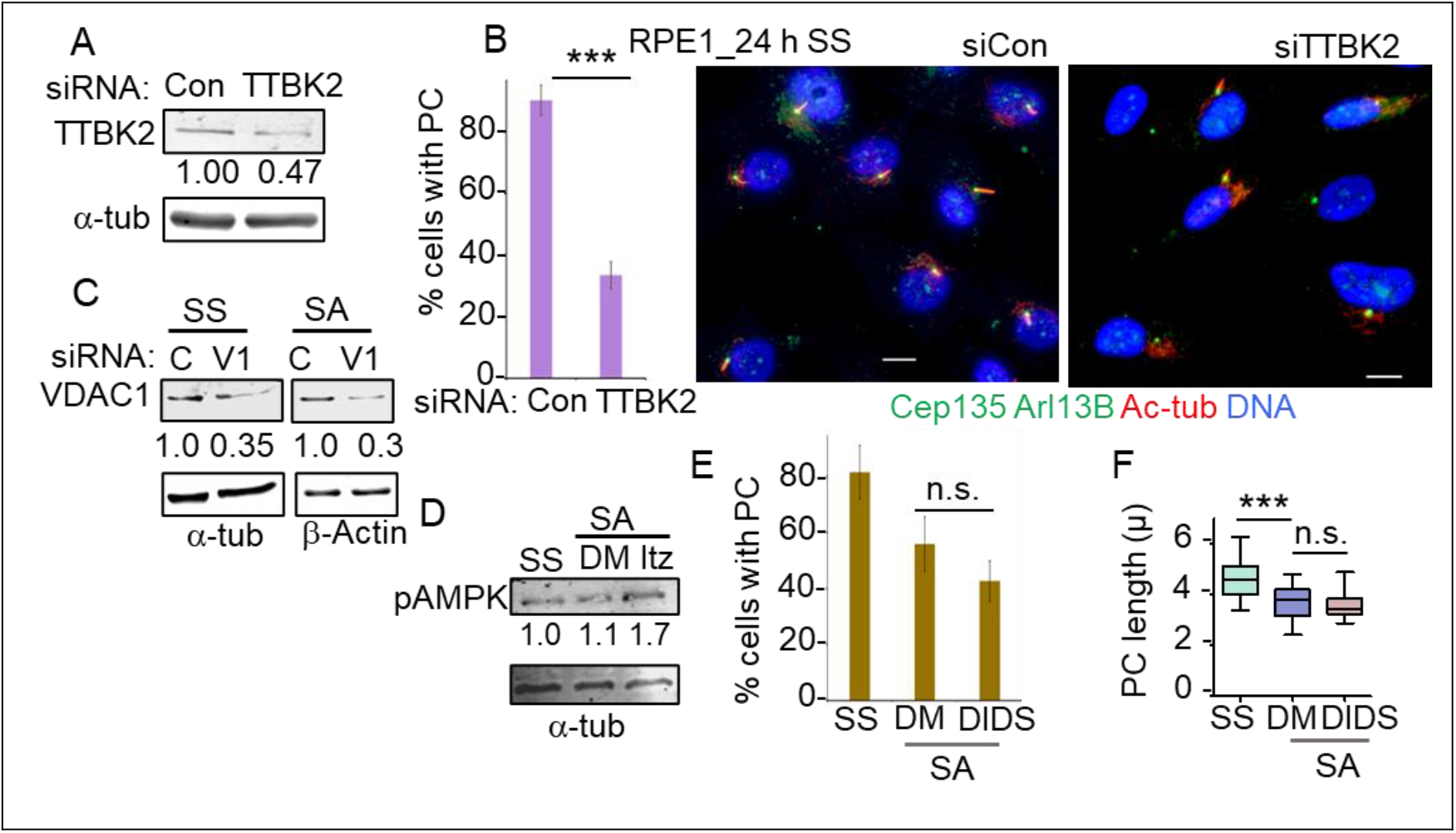
(A) RPE1 cells treated with the control and TTBK2-specific siRNAs, were serum-starved for 24 h. Representative immunoblots show the depletion of TTBK2 in siTTBK2-treated cells, where values show normalized intensities of TTBK2 band using α-tub as loading control. (B) Those cells were analyzed for the presence of PC, and plotted as bars where values represent mean ± S.D., n=3, at least 500 cells were counted per replicate. Representative micrographs of random fields of those cells are shown, where bar = 5 μ. (C) Representative immunoblots show VDAC1 level in siVDAC1#1-treated (V1) RPE1 cells collected during SS and SA condition. Values show normalized intensities of VDAC1 bands, where α-tub or β-actin was used as loading control. (D) RPE1 cells were starved (SS) and then serum was added (SA) to those cells following the strategy in Fig.4A. Itraconazole (Itz) was added during SA as in Fig.4F. Representative immunoblots show the level of phospho-AMPK (pAMPK) the indicated cells, where values show normalized intensities of phospho-AMPK, using α-tub as loading control. (E-F) In a similar experiment, DIDS was added during SA. Cells stained for PC markers were analyzed for percent ciliation (E; n=3, at least 500 cells were counted per replicate) and PC length (F; n=40-50 cells per replicate), in a similar manner as described earlier. DMSO (DM) was used as solvent control.

